# Investigation of immune response to Mesenchymal Stromal Cell-derived Extracellular Vesicles in the cancer setting

**DOI:** 10.1101/2025.01.28.635343

**Authors:** Katie E Gilligan, Oliver Treacy, Clodagh P O ’Neill, Elan C McCarthy, Emma McDermott, Elma Mammen, Peter Dockery, Aideen E Ryan, Róisín M Dwyer

## Abstract

Mesenchymal Stromal Cell derived extracellular vesicles (MSC-EVs) may retain the cancer targeting and immune privilege of MSCs. The immense potential MSC-EVs hold as tumour-targeted therapeutics warrants an understanding of potential adverse events to support clinical translation. This study aimed to determine whether MSC-EVs would elicit an immune response following administration in tumour-bearing immunocompetent animals. Secreted EVs were isolated from both human and murine bone marrow derived MSCs and characterized. hMSC-EVs or mMSC-EVs were administered intravenously into 4T1 breast tumour-bearing Balb/c mice or healthy controls. Tumour tissue, draining lymph nodes and spleens were harvested, dissociated into a single cell suspension and flow cytometry performed targeting T cells, myeloid derived suppressor cells (MDSCs), macrophages, dendritic cells and natural killer (NK) cells. The 4T1 model immune profile was first determined by comparing the spleen of tumour-bearing animals to healthy controls. T cells were increased in tumour-bearing animals (CD4+/CD25+ p=0.041; CD8+/CD25+ p=0.02). A significant elevation of GR-1+ MDSCs (p=0.002), CD11b+ macrophages (p=0.023) and CD11c+ dendritic cells (p=0.001) was also observed. In contrast, CD27+ NK cells were significantly decreased compared to the spleen of healthy animals (p=0.006). Collectively this data validated the immune profile and supported the determination of any changes in response to hMSC-EVs or mMSC-EVs administration. No significant activation of CD4+ (p=0.20) or CD8+ (p=0.57) T cells were seen in tumour tissue in both groups. The percentage of GR-1+ MDSCs (28% vs 27%, p=0.92), CD11b+, CD11c+ and CD27+ cells were similar regardless of EV origin. No significant changes in T cells, MDSCs, macrophages, dendritic or NK cells were observed in the lymph node or spleen of animals that received hMSC-EV versus mMSC-EVs. In conclusion, human MSC-EVs elicited no discernible immune response in mice, supporting the hypothesis that MSC-EVs retain the immune privilege of the secretory cell. This reinforces the therapeutic potential of MSC-EVs.

## 1. Introduction

In 2022, 2.3 million women were diagnosed with breast cancer, with 670,000 deaths recorded globally [1]. When diagnosed at an early stage, patients with breast cancer have an excellent prognosis. However, advanced metastatic breast cancer remains incurable, with limited treatment options available. Novel therapeutic strategies capable of targeting metastatic disease are urgently required. The goal for cancer treatment is the direct delivery of therapeutics to the site of metastatic disease while sparing healthy tissue. Mesenchymal Stromal Cells (MSCs) initially presented exciting potential in this arena due to a unique ability to home to the site of tumours and metastases when administered into the circulation, and to bypass immune recognition [2,3]. These cells play a critical role in wound healing and have the ability to differentiate into multiple lineages including bone, fat and cartilage. However, there are safety issues relating to MSC administration due to the inherent immune privilege and regenerative capacity of the cells. This raises the potential for masking of the tumour from immune detection, the secretion of chemokines and growth factors that could encourage tumour progression or recurrence, and the potential for cell transformation [4,5].

Small Extracellular Vesicles (sEVs) are released by all cells and are believed to retain features of the secreting cell. The lipid bilayer characteristic of sEVs protects a cargo of nucleic acids (mRNA, microRNA etc) and proteins that are shuttled between cells and throughout the circulation. Therefore MSC-sEVs hold promising potential as the vesicles may retain MSC tumour immune privilege and homing properties, while avoiding potential challenges associated with the use of cells [4-7]. Another potential benefit of MSC-sEVs is the expression of CD47 which binds to signal regulatory protein alpha (SIRP-α) and in turn initiates a “don ‘t eat me” signal. The signal inhibits macrophage phagocytosis, which has been reported to suppress sEV clearance, although further investigation is required [8].

sEVs present a variety of advantages including small size (<200nm) and biocompatibility, increased stability for encapsulation of therapeutic agents, potential for surface modification to improve homing, and the ability to fuse with the plasma membrane to support direct delivery of therapy [9]. Compared to cells, MSC-sEVs also present reduced safety issues surrounding systemic administration such as lack of tissue entrapment issues, and also potential for higher yield production [10]. Due to the rapid expansion of sEV research there has been a major focus on defining sEV subpopulations and establishing guidelines for isolation and characterisation. Methodology for modification of cargo and scale up of reproducible sEV production also continue to develop, all of which are crucial for clinical translation [11,12].

In the cancer setting, sEVs (both wild type and engineered) containing a range of therapeutic cargo have been employed preclinically as a means of therapy. In these models, most studies to date have used immunocompromised animals that preclude investigation of the immune response to the administered EVs [13-16]. Although sEVs have been extensively studied as potential therapeutics for immune-related diseases, their interaction with the host immune system remains poorly understood [17]. If a cell triggers an immune response, it is likely that the EVs it secretes would elicit a similar immune response, as EVs are thought to reflect the characteristics of the cell from which they originate [6].

There is a need to use human sEVs for therapeutic administration in In Vivo models, to allow future studies to be as clinically relevant as possible. MSC-sEVs have been administered in a clinical trial, not in the cancer setting, but for Graft vs Host Disease (GvHD) after the failure of standard treatments [18]. Improved clinical symptoms were observed with no negative side effects, and levels of Interleukin (IL)-1β, Tumour Necrosis Factor (TNF)-α and Interferon (IFN)-γ remaining stable throughout [18]. In a clinical trial for acute and chronic kidney disease, a larger study that included analysis of immune impact employed MSC-sEVs, and levels of TGF-β1, IL-10 and TNF-α were monitored with no significant changes observed when compared with the control group [19]. The safety and tolerance of MSC-sEVs observed in these studies is promising in the clinical setting, however, in cancer very little research has focused on the immune implications of EV administration [15]. Emerging clinical trials underscore the growing potential of EV-based therapies for cancer e.g. Codiak Bioscience Phase I trial of exoSTING (NCT04592484) explores EVs activating the STimulator of InterferoN Genes (STING) pathway, while another Phase I trial (NCT03608631) evaluates MSC-derived exosomes loaded with KRASG12D siRNA (iExosomes) for treatment of metastatic pancreatic cancer. These studies highlight the promise of EV-based therapies while emphasizing the critical need to investigate host-EV interactions to enable successful clinical translation.

The tumour microenvironment is complex and variable comprising of a range of cell types including immune cells which contribute to tumour progression. The tumour can exploit the immune system to aid progression by recruiting cells to the site to mask detection by the host [20]. Immunotherapies have been developed to target such tumours but this can depend on tumour site and stage of the disease [21]. In a previous study by our group, human MSC-sEVs encapsulating miR-379 showed promising therapeutic impact after administration in an immune compromised model of aggressive breast cancer [22]. Given the promising results of that and other studies, and progress of MSC-sEVs towards clinical trials for cancer, it is important to determine whether there will be an immune response to MSC-sEVs in the cancer setting. This study therefore aimed to successfully isolate sEVs from both murine and human MSCs to investigate the host immune response after administration of these MSC-sEVs in the presence of breast cancer.

## 2. Materials and Methods

### 2.1 Ethics statement

Preclinical studies were performed following approval by the Institutional Animal Care Research Ethics Committee (ACREC Ref. 18-Jun-02) at University of Galway and a licence (AE19125/P084) from the Health Products Regulatory Authority (HPRA). The HPRA has the authority in Ireland for implementation of EU legislation (Directive 2010/63/EU).

### 2.2 Isolation and Culture of Mesenchymal Stromal Cells

As described in a previous publication [22] adult human bone marrow (iliac crest) MSCs (hMSCs) were isolated from healthy volunteers using a defined clinical protocol by collaborators in the Regenerative Medicine Institute at University of Galway (Research Ethics Committee Ref. 08/May/14). Murine MSCs (mMSCs) were also isolated from the bone marrow of Balb/c mice as previously described [23]. The femur and tibia were removed from mice after euthanising by CO_2_ inhalation, and placed in MEM-α supplemented with 10% heat-inactivated fetal bovine serum (FBS), 10% equine serum, and 1% penicillin/streptomycin. Using a 30.5G needle, cells were flushed from the epiphysis of each bone with culture medium and filtered through a 70-μm cell strainer to remove clumps. Cells were then centrifuged at 400 × g for 5 min, re-suspended in 25 mL of culture medium, and plated. MSCs from both sources were cultured in MEM-α supplemented with 5% serotyped FBS, 100UI/mL Penicillin/100μg/mL Streptomycin and basic human Fibroblast Growth Factor (FGF) (4ng/ml).

### 2.3 EV Isolation

To prepare EV-depleted media, MEM-α media supplemented with 20% serotyped FBS was filtered (0.22μm), followed by ultracentrifugation (Hitachi himac micro ultracentrifuge CS 150fNX, S50A-2152 rotor) at 110,000 x g for 18 hours at 4°C. The final media was then diluted 1:1 in basal MEM-α. mMSCs or hMSCs were seeded at 2 x 10^6^ cells/T175 cm^2^ flask and cultured in EV-depleted media for 48hrs, prior to isolation. EVs were isolated from the cell conditioned media of 12 x T175 cm^2^ flasks, through a series of differential centrifugation steps (300 x g for 10 mins followed by 2000 x g for 10 mins), followed by microfiltration (0.22μm) and two sequential ultracentrifugation steps at 110,000 x g for 70 minutes each. The isolated murine and human MSC-EVs were placed in aliquots for In Vivo administration (PBS), Nanoparticle Tracking Analysis (NTA) (PBS) or protein analysis (lysis buffer including protease inhibitors and sodium orthovanadate) and stored at -80°C until required for application. Prior to In Vivo administration, isolated EVs were pooled to support administration of 1 x 10^8^ in 100μl.

### 2.4 Nanoparticle Tracking Analysis

Nanoparticle tracking analysis (NTA, NanoSight NS500, Malvern Panalytical, UK) was employed to obtain EV particle size distribution and concentration as previously described using a 405 nm laser source and EMCCD camera. NTA software version 3.2 was run using optimised and validated protocols [24]. NS500 calibration was verified daily using 100 nm polystyrene latex calibration nanoparticles (Malvern Panalytical). A total of five 60 second videos were recorded for each sample.

### 2.5 Western Blot

To determine protein concentration the MicroBCA Assay (Pierce™, Thermo Fisher Scientific, Massachusetts, USA) was used according to manufacturer ‘s instructions. Western blot was performed targeting EV associated CD63 (abcam: ab59479), CD81 (abcam; ab79559), endosomal protein TSG101 (abcam; ab125011) and negative marker Calnexin (abcam; ab22595). Briefly, 10μg of protein samples were denatured at 70°C for 10 min. Samples were loaded on a pre-cast Mini-PROTEAN® TGX™ Gel (Bio-Rad) and run for 40 min at 200 V. Protein molecular weight standards (MagicMark, Invitrogen 20–220 kDa) were run simultaneously on each gel. Transfer to a nitrocellulose membrane was performed and blots were blocked in 5% milk in Tris-Buffer saline (TBS-T) for 2 h. All antibodies were diluted in 0.1% milk in TBS-T at 1:1000 dilution and each membrane was incubated with CD81 (1.5 h, RT), or CD63, TSG101 or Calnexin overnight at 4°C. Following a series of washing steps, a solution of secondary antibody was added [CD81—1:10,000 rabbit anti-mouse IgG HRP; Abcam ab6728; CD63, TSG101, Calnexin -all 1:3000 goat anti-rabbit IgG HRP; Abcam ab6721] to each membrane for 1.5 hours. In order to visualize the membranes using the Gel Doc™ XR+ and ChemiDoc™ XRS + Systems, clarity™ Western ECL (Bio-Rad Laboratories, Maryland, USA) chemiluminescent substrate solution was applied to each membrane and analyzed using Image Lab™ Software (Bio-Rad, version 5.2.1).

### 2.6 Transmission Electron Microscopy

Isolated EVs were fixed by embedding in resin as previously described [25]. Briefly, prior to ultracentrifugation, a primary fixative was added to each sample (2% glutaraldehyde, 2% paraformaldehyde in a 0.1 M sodium cacodylate/HCL buffer, pH 7.2). EVs were then submerged in a secondary fixative (1% osmium tetroxide), and dehydrated in a series of alcohol washes, before being embedded in resin (Agar Scientific commercial kit). The EVs in resin were sectioned and loaded onto a copper grid, stained using Uranyl Acetate, and viewed using a Hitachi H7000 transmission electron microscope.

### 2.7 Analysis of EV administration in an immune competent In Vivo model

Eighteen female Balb/c mice aged 6-8 weeks (Charles River Laboratories Ltd. Kent, UK) were used in the study. Twelve mice under anesthesia received a 4^th^ inguinal mammary fat pad injection of 1 x 10^5^ 4T1 cells (ATCC-CRL-2539, cultured in RPMI, 10% FBS and 1% P/S) suspended in 200μl of RPMI and tumours were allowed to form (study timeline shown in Fig. 1 A). Cells were injected under sterile conditions on a surgical mount/heated stage. All animals were drug naïve, with no previous procedures performed, and weighed between 17–20g. Weight and general behaviour were monitored daily and scored. The animal ‘s environmental enrichment was monitored daily, including cage conditions, food and water, health and behaviour. Tumour progression was monitored by visual inspection and regular caliper measurements. Fifteen days following tumour induction and once the tumours were palpable, animals received an IV injection of 1x 10^8^ hMSC-EVs (n=3) or mMSC-EVs (n=3), with n=6 tumour bearing animals receiving no EV dose. Similarly, tumour free animals (n=9) received 1 x 10^8^ hMSC-EVs (n=3) or mMSC-EVs (n=3) or remained EV free (n=3). Three days later, animals were sacrificed and tumour tissues, draining lymph nodes (LN) and spleens were collected for digestion and flow cytometry analysis.

**Figure 1.**
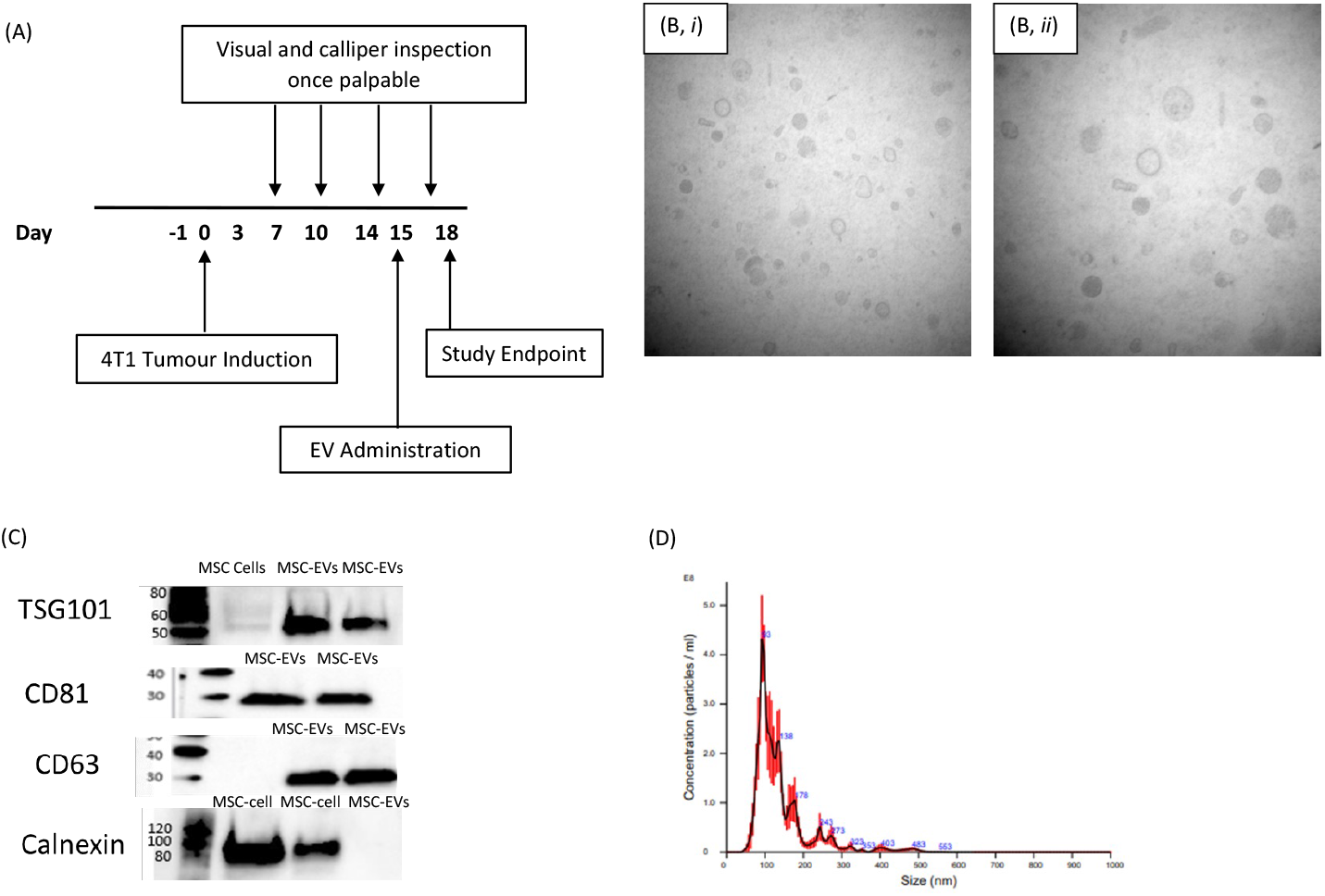
In Vivo timeline and EV characterisation. (A) Timeline of the In Vivo study with tumour induction performed at day 0 and EV (1 x 10^8^) tail vein administration performed on day 15. (B) TEM imaging of the MSC-EVs embedded in resin and visualised at both (i) wide field (50,000 X) and (ii) close field view (80,000 X) showing the distinctive lipid bilayer. (C) Western blot analysis of the MSC cells and corresponding secreted EVs – positive for endosomal marker TSG101, tetraspanins CD63 and CD81 and negative for calnexin [full-face blots available in Supplementary Figure 2]. (D) NTA analysis determined the concentration and size of the EVs as <200nm indicating small EVs. The red shading indicates the range of readings across the 5 replicates performed.

### 2.8 Flow cytometry analysis

Tumour tissues were digested using the Miltenyi Biotec Tumor Dissociation Kit. Tumour, draining lymph nodes and spleens were pushed through a cell strainer (40μm Nylon Cell Strainer, BD Falcon) to dissociate into PBS, centrifuged at 400 x g for 5mins and resuspended in a single cell suspension in FACS buffer (PBS and 1% FBS). Cells were seeded into a 96 well plate at 100,000 cells/well in triplicate. Appropriate antibodies (all MSC Biolegend; MSC Dublin, Ireland; see Table 1 for details) were added to each of the wells and incubated for 30 mins before resuspension in FACS buffer.

**Table 1.** Antibodies used for flow cytometry.

| **Panel 1 Antibodies (Clone)** | **Catalog no.** | **Panel 2 Antibodies (Clone)** | **Catalog no.** |
| --- | --- | --- | --- |
| CD4-APC (GK1.5) | 100412 | CD11b-FITC (DX5) | 108910 |
| CD8-FITC (53-5.8) | 140403 | CD11c-FITC (N418) | 117306 |
| GR-1-PerCP-Cy5 (RB6-8C5) | 128001 | MHC-I-APC-Cy7 (SF1-1.1) | 116629 |
| Ly-6c-PE-Cy7 (HK1.4) | 108415 | MHC-II-APC (39-10-8) | 115003 |
| CD19-BV510 (6D5) | 115545 | CD27-BV510 (LG.3A10) | 124229 |

Abbreviations – APC: Allophycocyanin; PE: Phycoerythrin, PerCP: Peridinin Chlorophyll Protein Complex; BV: Brilliant Violet; FITC: Fluorescein isothiocyanate; CD: Cluster of differentiation; GR-1: glucocorticoid receptor; MHC: major histocompatibility complex.

Samples were acquired using the BD FACSCanto II flow cytometer and analysed using FlowJo analysis software version 10 (Tree Star, Ashland, OR, USA). Fluorescence minus one (FMO) controls were used to position the gate to determine the percentage of live cells that stained positive for the target marker. Gating strategies are provided in Supplementary Figure 1. Single stain controls were used to compensate for spectral overlap (Table 1). Statistical analysis was performed using GraphPad Prism version 9.2.0. Each cell suspension and antibody combination from every individual tissue was tested in triplicate wells and the percentage of live cells positive for the marker of interest determined. The mean value of three readings from each sample representing an individual tissue from each animal was then calculated, and the mean values across animal groups compared using a two-tailed paired t test.

## 3. Results

### 3.1. EV Characterisation

EVs were isolated from hMSC and mMSC conditioned media and characterised by western blot, TEM and NTA (Figure 1 B-D). EVs embedded in resin were visualised under a transmission electron microscope (Figure 1 B). Widefield (50,000 X) view showed a large number of EVs of the correct size (50-120nm) and shape (Figure 1 B, i). Closefield (80,000 X) view showed the characteristic lipid bilayer of the vesicles (Figure 1 B, ii). Western blot analysis revealed the presence of Tetraspanins CD63 (25-30kDa), CD81 (26-30kDa), the endosomal marker TSG101 (50-60kDa) and the absence of endoplasmic reticulum marker Calnexin (80-90kDa) (Figure 1 C). NTA results showed particle sizes below 200nm. Clear peaks were seen between the 30-150 nm size showing sEVs across all five readings (Figure 1 D), with concentrations ranging from 4.08 x 10^9^ - 6.6 x 10^9^ particles/ml.

### 3.2. Impact of tumour formation (EV free): Host immune activation in spleen of 4T1 tumour bearing animals compared to healthy controls

The immune microenvironment in tumours is shaped by the tumour mutational landscape, vasculature, expression of cell surface and secreted immunomodulatory mechanisms. The presence of CD4^+^ and CD8^+^ T cells within the tumour have been linked to overall survival in cancer patients. However, the tumour microenvironment is antagonistic to T cells and depletion is observed as the cancer progresses [26,27]. When the spleens of the tumour bearing animals were compared to the non-tumour bearing there were several significant changes in immune cell populations observed. In this study, spleen CD4^+^ T cells were significantly decreased (p<0.001) when compared to the non-tumour bearing animals, while the subset of activated CD4^+^/CD25^+^ cells was increased (Figure 2 A i, ii). CD8^+^ T cell levels remained unchanged between the groups (11.5% vs 12%; p=0.49; Figure 2 A, iii). No change was observed in the frequency of CD8+ cells, however similar to CD4+ cells, the frequency of activated CD8+ T cells as determined by the expression of CD25 was significantly increased in the presence of cancer (2.4% vs 8.8%; p=0.02, Figure 2 A; iv). MDSCs including GR-1^+^ cells play an immunosuppressive role in the tumour microenvironment [28,29], and levels were significantly increased in the spleen of tumour bearing animals (p=0.0016). The subset of GR-1^+^/Ly-6C^+^ positive cells was significantly decreased in tumour bearing animals (MFI-12161 vs 16307; p=0.02 Fig.2 B i, ii). Levels of CD11b^+^ macrophages were significantly increased (7.2% vs 19.7%; p=0.02, Figure 2 C, i). Subset markers of the CD11b^+^/MHCI^+^ were decreased in the tumour bearing group (MFI-8068 vs 12507; p=0.03) with CD11b^+^/MHCII^+^ cells increased (Figure 2 C, ii, iii; p<0.001)). CD11c^+^ dendritic cells were significantly increased in the spleen of tumour bearing animals compared to the non-tumour bearing mice (p=0.0006, Figure 2 D, i). CD11c^+^/MHCI^+^ (MFI-16354 vs 19786; p=0.03) and CD11c^+^/MHCII^+^ (MFI-896 vs 1629; p=0.03) were both decreased in the tumour bearing group (Fig.2 D, ii, iii). CD27^+^NK cells are important for recognising and eradicating cancer cells, and levels were significantly decreased in the spleen of tumour bearing animals compared to the non-tumour bearing (p=0.0058; Figure 2 E). CD19^+^ B-cells were activated at similar levels regardless of tumour presence (p=0.22; Figure 2 F). Overall, the tumour profile showed activation of T cells and recruitment of MDSCs in parallel with decreasing NK cells.

**Figure 2.**
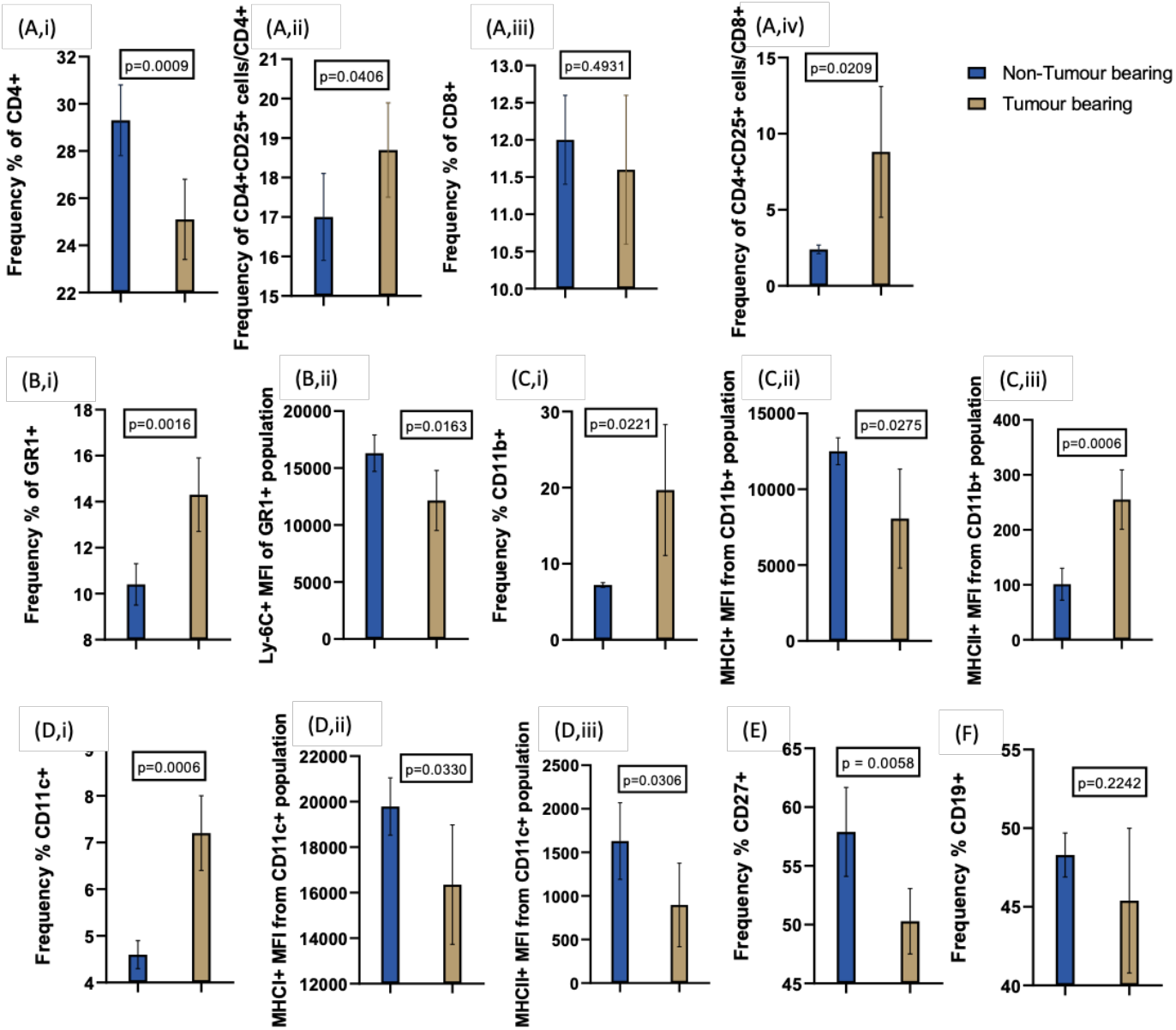
Host immune activation in spleen of 4T1 tumour bearing animals compared to non-tumour bearing. (A) T cell activation – (A, i) CD4^+^ T helper cells were significantly decreased (25% vs 29%; p< 0.001) when compared to the non-tumour bearing mice. (A, ii) The subset marker of activation CD4^+^/CD25^+^ was significantly increased in the tumour bearing animals (17% vs 18.7%; p=0.04). (A, iii) CD8^+^ T cell levels remained unchanged (11% vs 12%; p=0.49). (A, iv) CD8+/CD25+ cells were significantly increased in the tumour bearing animals (2.4% vs 8.8%; p=0.02). (B, i) GR-1^+^ levels were significantly increased in the tumour bearing animals (10.4% vs 14.3%; p=0.00). (B, ii) The subset of GR-1^+^/Ly-6C^+^ (MFI-12161 vs 16307; p=0.01) positive cells was significantly decreased in tumour bearing animals. (C, i) Levels of CD11b^+^ macrophages were significantly increased in the tumour bearing animals (7.2% vs 19.7%; p=0.02). (C, ii) Subset markers of the CD11b^+^/MHCI^+^ were decreased in the tumour bearing group (MFI-8068 vs 12507; p=0.02) with (C, iii) CD11b^+^/MHCII^+^ cells increased (MFI-255 vs 101; p=0.00). (D, i) CD11c^+^ dendritic cells were significantly increased in the spleen of tumour bearing animals compared to the non-tumour bearing mice (4.6% vs 7.2%; p=0.00). (D, ii) CD11c^+^/MHCI^+^ (MFI-16354 vs 19786; p=0.03) and (D, iii) CD11c^+^/MHCII^+^ (MFI-896 vs 1629; p=0.03) were both significantly decreased in the tumour bearing group. (E) CD27^+^ levels (57.9% vs 50.3%; p<0.006 were significantly decreased in the tumour bearing animals compared to the non-tumour bearing. (F) B-cells were activated at similar levels regardless of tumour presence (48.3% vs 45.4%; p=0.22).

### 3.3 Host immune activation in 4T1 tumour following IV administration of either human or murine MSC-derived EVs

EVs isolated from human or murine MSCs were administered to animal bearing 4T1 tumours and non-tumour bearing animals. This was to determine the impact on the host immune system with and without the presence of a tumour. The tumour single cell suspension was probed for markers including T cells, MDSCs, NK cells and B cells to detect an infiltrating host immune response to the administered EVs. No significant activation of T cells was seen in either the group that received murine MSC-EVs or human MSC-EVs. This was indicated by similar frequencies of CD4^+^ and CD8^+^ populations (Fig. 3 A, i, iii). CD25^+^ subsets indicating activation of CD4^+^ and CD8^+^ cells also remained unchanged (Fig. 3 A ii, iv), in either murine or human MSC-EV treated tumours. MDSCs and dendritic cell activation was similar between both groups regardless of EV origin (Fig. 3). CD27^+^ NK cells were detected at similar levels as were B-cells. Overall, no significant changes we observed in any of the innate immune cell populations following human or murine MSC-EV administration.

**Figure 3.**
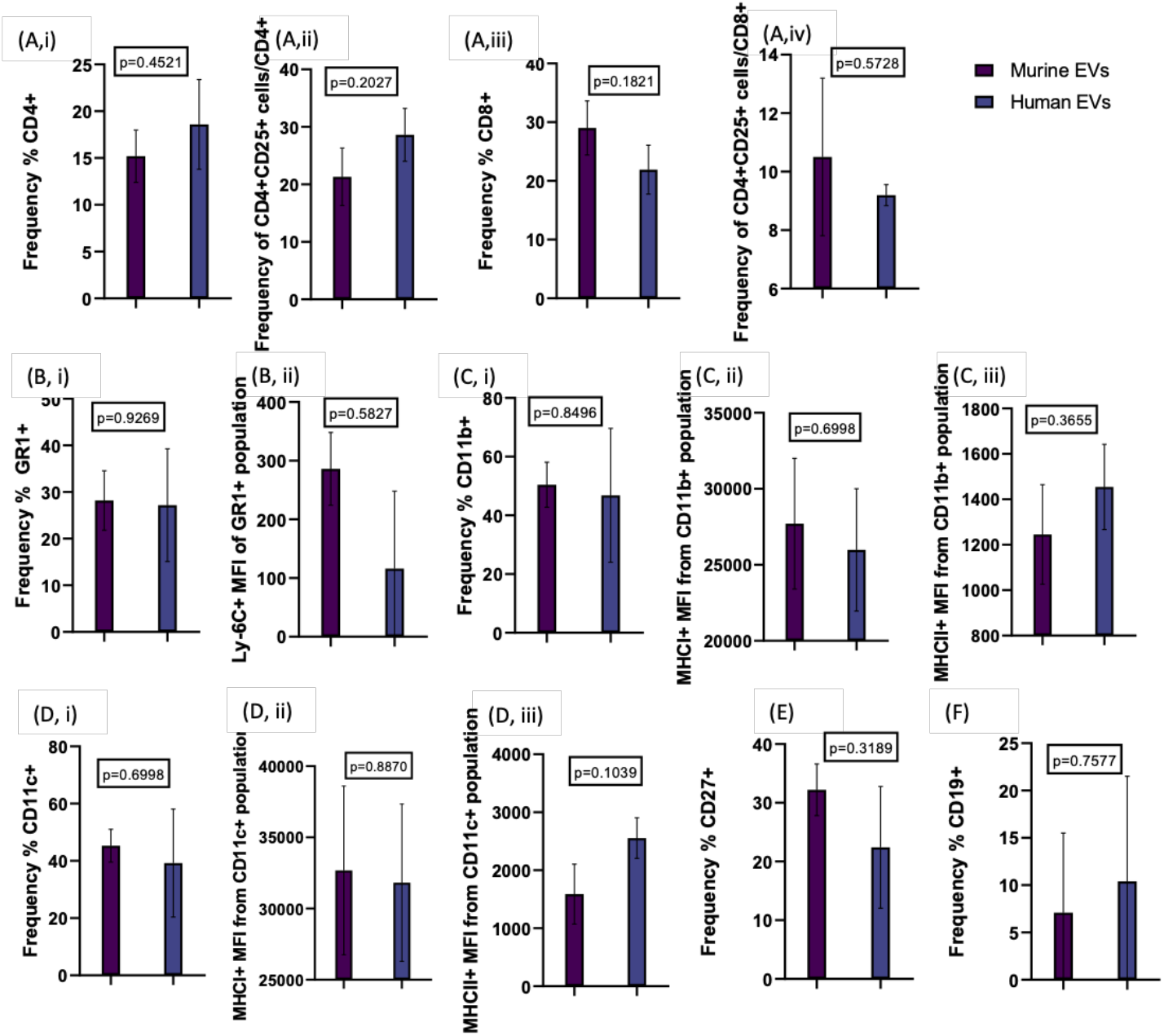
Host immune activation in cell suspension of a 4T1 tumour following IV administration of either human or murine MSC-derived EVs. (A, i) Levels of CD4+ (15.2% vs 18.6%, p=0.45) and (A, ii) CD25+ subsets indicating activation of T helper cells (21.3% vs 28.6%; p=0.20) remained unchanged. (A, iii) CD8+ (21.3% vs 28.6%, p=0.18) populations remained stable with (A, iv) CD25+ subsets indicating activation of cytotoxic T cells (29% vs 21.9%; p=0.57) were not significantly elevated in either group. (B, i) Levels of GR-1+ (28.2% vs 27.2%, p=0.92) and (B, ii) GR-1+/Ly-6c+ (MFI-286 vs 116, p=0.58) remained stable. (C, i) CD11b+ (50.4% vs 46.8%, p=0.85), along with the subset markers (C, ii) CD11b+/MHCI+ (MFI-27707 vs 25979; p=0.70) and (C, iii) CD11b+/MHCII+ (MFI-1245 vs 1454; p=0.37) remained consistent. (D, i) CD11c+ showed no significant change (45.3% vs 39.2%; p=0.70) which was also true of the subset markers (D, ii) CD11c+/MHCI+ (MFI-32681 vs 31813; p=0.89) or (D, iii) CD11c+/MHCII+ (MFI-1589 vs 2554; p=0.10). (E) CD27+ NK cells were detected at similar levels (32.2% vs 22.4%; p=0.32) as were (F) CD19+ positive cells (7.1% vs 10.4%; p=0.76).

### 3.4 Host immune activation in the lymph nodes of 4T1 tumour bearing or non-tumour bearing animals following IV administration of either human or murine MSC-derived EVs

When the lymph nodes were assessed no significant changes in activation of any immune cell populations were observed between the animals receiving murine EVs when compared to the animals receiving human EVs. This was indicated by stable activation of CD4^+^ (54.3% vs 51.1%, p = 0.68; Figure 4 A i) and CD8^+^ T cells (19% vs 18%, p = 0.58, Figure 4 A iii), along with CD25^+^ subsets of each (Figure 4). There was no significant activation of MDSCs as GR-1^+^ (4.8% vs 2.0%, p=0.50) and GR-1^+^/Ly-6C^+^ (MFI-54667 vs 68020, p=0.27) showed similar activation regardless of EV origin (Figure 4 B i, ii). CD11b^+^ macrophages (10.0% vs 5.7%; p=0.55) and subsets of CD11b^+^/MHCI^+^ (MFI-13241 vs 14255; p=0.62), CD11b^+^/MHCII^+^ (MFI-100 vs 136; p=0.17) were all activated at similar levels between EV groups (Figure 4 C i, ii, iii). NK cell levels and B cells were also not affected by source of EVs administered (Figure 4). Lower cell number in the LN meant that not all targets could be detected, however no changes were observed in any of the cell populations that were analysed successfully.

**Figure 4.**
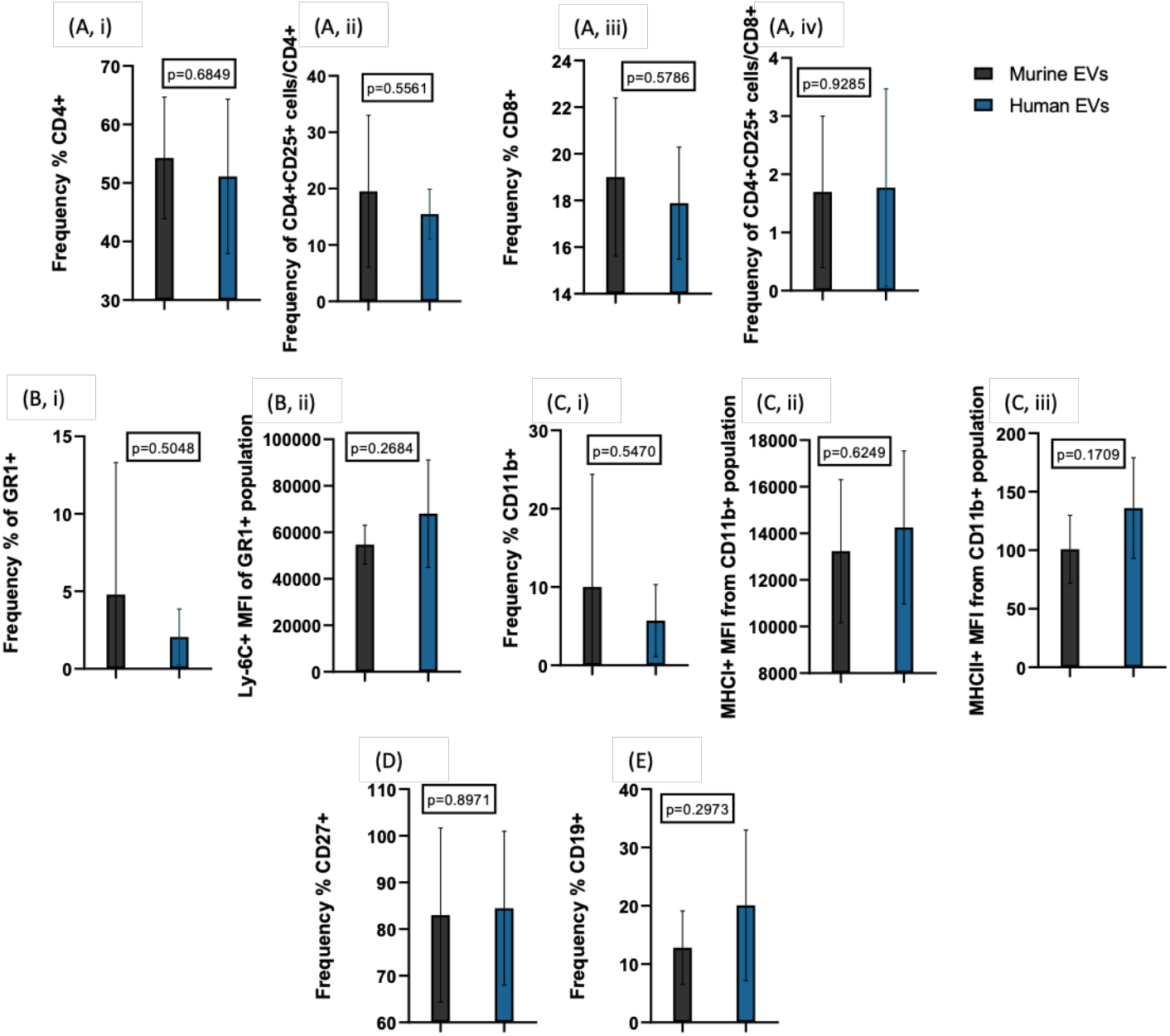
Host immune activation in the lymph nodes of 4T1 tumour bearing or non-tumour bearing animals following IV administration of either human or murine MSC-derived EVs. Stable activation of T helper or cytotoxic T cells, as (A, i) CD4+ (54.3% vs 51.1%, p = 0.68), (A, ii) CD4+/CD25+ (19.5% vs 15.4%; p=0.56 and (A, iii) CD8+ (19% vs 17.8%, p = 0.58), (A, iv) CD8+/CD25+ (1.7% vs 1.77%; p=0.93) were observed. There was no significant activation of (B, i) GR-1+ (4.8% vs 2.0%, p=0.50) and (B, ii) GR-1+/Ly-6C+ (MFI-54667 vs 68020, p=0.27). (C, i) CD11b+ (10.0% vs 5.7%; p=0.55) and subsets (C, ii) CD11b+/MHCI+ (MFI-13241 vs 14255; p=0.62), (C, iii) CD11b+/MHCII+ (MFI-100 vs 136; p=0.17) were all activated at similar levels between EV groups. (D) CD27+ cell levels and (E) CD19+ were also not affected by source of EVs administered (83% vs 84.4%; p=0.90; 12.8% vs 20%; p=0.30).

### 3.5 Host immune activation in the spleen of 4T1 tumour bearing or non-tumour bearing animals following IV administration of either human or murine MSC-derived EVs

The spleens of animals were also analysed, T cells were activated at similar levels, showing no significant changes in CD4^+^ (Figure 5 A i - 27% vs 27% p=0.97), CD4^+^/CD25^+^ (Figure 5 A ii - 18.4% vs 17.3%; p=0.23) or CD8^+^ (Fig. 5 A iii - 12.2% vs 11.4%, p=0.17), CD8^+^/CD25^+^ (Fig. 5 A iv - 6.5% vs 4.7%; p=0.52) populations. MDSCs were also expressed at similar levels with no changes seen between groups as indicated by GR-1^+^ (12.7% vs 12.0%; p=0.65) and GR-1^+^/Ly-6C^+^ (p=0.36) levels (Figure 5 B i, ii). Macrophages remained stably activated regardless of human or murine EV administration as seen by CD11b^+^ (p=0.68), CD11b^+^/MHCI^+^ (p=0.72), and CD11b^+^/MHCII^+^ (p=0.92, Figure 5 C i, ii, iii). CD11c^+^ dendritic cells were unchanged in animals that received either murine or human EVs (Figure 5 D). No changes were observed in NK cells or B cells in response to EV source (Figure 5 E, F).

**Figure 5.**
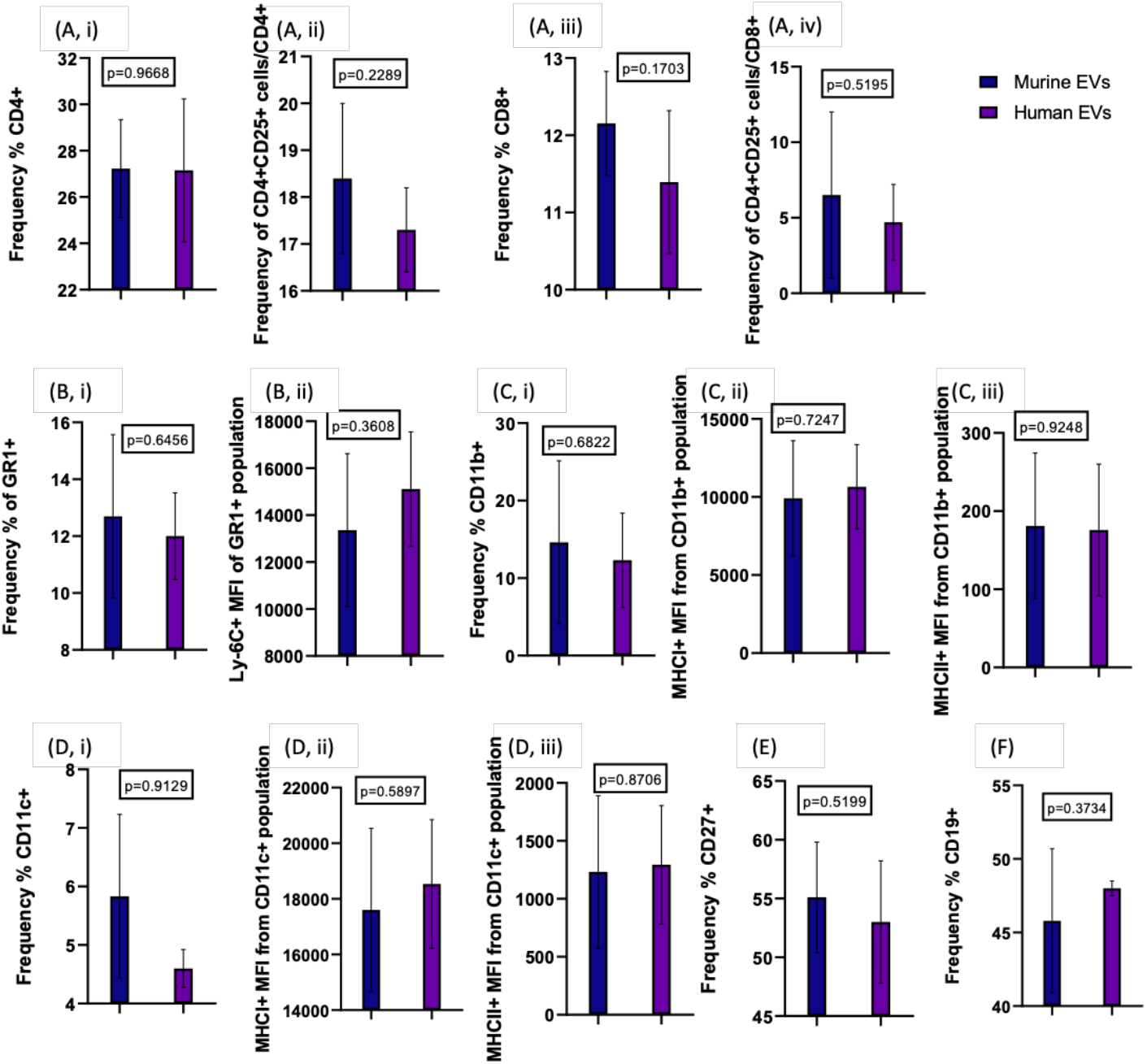
Host immune activation in the spleen of 4T1 tumour bearing or non-tumour bearing animals following IV administration of either human or murine MSC-derived EVs. T cells (both helper and cytotoxic) were activated at similar levels, showing no significant changes in (A, i) CD4+ (27% vs 27% p=0.97), (A, ii) CD4+/CD25+ (18.4% vs 17.3%; p=0.23) or (A, iii) CD8+ (12.2% vs 11.4%, p=0.17), (A, iv) CD8+/CD25+ (6.5% vs 4.7%; p=0.52) populations. (B, i) GR-1+ (12.7% vs 12.0%; p=0.65) and (B, ii) GR-1+/Ly-6C+ (MFI-13359 vs 15109; p=0.36 levels were not significant. (C, i) CD11b+ remained stably activated regardless of human or murine EV administration (14.6% vs 12.3%; p=0.68), (C, ii) CD11b+/MHCI+ (MFI-9915 vs 10659; p=0.72), (C, iii) CD11b+/MHCII+ (MFI-181 vs 175; p=0.92). (D, i) CD11c+ was unchanged (5.8% vs 5.9%; p=0.91), (D, ii) CD11c+/MHCI+ (MFI-17604 vs 18536; p=0.59) and (D, iii) CD11c+/MHCII+ (MFI-1231 vs 1294; p=0.87). No changes were observed in (E) CD27+ (55.1% vs 53.0%; p=0.52) or (F) CD19+ (45.8% vs 48%; p=0.37) in response to EV source.

## 4. Discussion

For clinical translation in the cancer setting, it is critical that there is research focusing on the host immune response to EV administration. The impact of administration of both murine and human MSC-EVs on the host immune system with and without the presence of the tumour, was the focus of this study.

Immune competent animals received either human or murine EVs, and the tumours, lymph nodes and spleens of all animals were digested, and flow cytometry analysis was performed. A broad range of immune markers were selected for analysis covering T cells (CD4^+^, CD8^+^; ^+^/CD25^+^), myeloid-derived suppressor cells (MDSC-GR-1^+^ CD11b^+^), dendritic cells (CD11c^+^), B cells (CD19^+^) and NK cells (CD27^+^). The impact of the 4T1 tumour model on the murine immune system is well established [30]. 4T1 tumours have been found to suppress the host immune system in order to avoid immunosurveillance and allow for tumour progression [27-29,31-34]. Therefore, prior to analysis of the impact of MSC-EVs, this baseline was validated by comparing the spleens of animals bearing 4T1 tumours to non-tumour bearing controls. This revealed there was an activation of several cell populations including T cells (CD4^+^/CD25^+;^ CD8^+^/CD25^+^). Turbitt et al [27], as part of a study investigating the effect of physical activity on tumour progression also investigated immune cell markers in the spleen of Balb/c mice bearing 4T1 tumours. Animals bearing 4T1 tumours without any intervention were compared to healthy controls to develop baseline readings. At the two-week time point, in line with the data presented here, 20% of CD4^+^ T cells and 10% of CD8^+^ T cell positive cells were observed. The levels of both T cell populations decreased over the following three weeks, suggesting the tumour was exploiting the immune system to remain undetected. It was shown that levels of T cells decreased as levels of GR-1^+^/CD11b^+^ cells increased over a five week period of 4T1 tumour growth [27].

GR-1^+^ MDSCs in our study were significantly increased in the tumour bearing animals (10.4% vs 14.3%; p=0.0016), although the subset GR-1^+^/Ly-6c^+^ was decreased in the tumour bearing group, similar to literature showing Ly-6c^+^ decreases as other MDSC populations increase [35]. CD11b^+^ and CD11b^+^/MHCII^+^ were both significantly increased in the tumour bearing animals indicating that MDSC recruitment was beginning [27,29,33]. Breast cancer recruits MDSCs to the tumour microenvironment to downregulate anti-tumour immunity and aid in tumour progression, particularly in 4T1 models [28,32,34]. Trad et al [29] also reported that increased levels of CD11c^+^ in 4T1 tumours meant that dendritic cells are recruited to help suppress CD4^+^ and CD8^+^ T cells, observed by an increase in the levels of CD11c^+^. This correlates with the findings in our study where CD11c^+^ levels were seen to be significantly elevated in tumour bearing animals.

In the presence of breast cancer, natural killer (NK) cells have been found to be decreased by deregulation of janus kinases (JAK) and the signal transducer and activator of transcription (STAT) pathway. Inhibition of this pathway has been found to help in progression of metastasis by in-turn decreasing NK cell anti-tumour activity [31]. Significantly decreased levels of CD27^+^ NK cells were seen in this study in the tumour bearing animals. As a whole, this data validated the immune profile of the breast cancer model and supported the determination of any changes in response to administration of human or murine MSC-EVs.

When tumour bearing animals that received mMSC-EVs were compared to animals that received hMSC-EVs, no changes in the frequency of T cells, MDSCs, macrophages, dendritic or NK cells were observed in the draining lymph node or spleen of animals. Due to the small size of the LN there was a lower cell number received after single cell digestion, therefore not all markers were detected in this cell population. No changes in weight or adverse events were observed during the study. Taken together, these results support that the administration of human or murine EVs had no significant effect on the host immune response.

Mendt et al [36] administered MSC-EVs for the treatment of cancer in an immune competent model. In the pre-clinical study BJ Fibroblast EVs were engineered to carry an siRNA for the treatment of pancreatic cancer. While no immune analysis was performed on the tumour bearing animals receiving EVs, parallel administration of BJ-EVs, MSC-EVs, or MSC-iExosomes (electroporated to contain the siRNA to KrasG12D) over a 3-week period to non-tumour bearing animals was performed followed by immunophenotyping of the spleen, bone marrow and thymus. Lymphocytes (CD3+, CD4+, CD8+, and CD19+) along with myeloid cells (CD11b+, F4/80+) showed no significant changes in the animals that received EVs [36]. This has now moved forward into a Phase I clinical trial (NCT03608631). This trial recruitment phase began in January 2021 with an extended completion date in April 2025. Patients are to receive 3 courses of IV delivered iExosomes. The study will measure tolerated dose and dose limiting toxicities [36]. This trial will be a promising step forward for MSC-EVs in the cancer setting.

Codiak Biosciences have also recently completed a Phase I clinical trial study focusing on therapeutic EV administration for a range of solid cancers - exoSTING (NCT04592484). The EV therapy involves activation of the STimulator of InterferoN Genes (STING) pathway which resulted in tumour regression in preclinical models. Although not involving MSC-EVs, it does demonstrate how the field is moving forward [37]. The preclinical study did not investigate the host immune response to administration of HEK293 EVs in the immune competent model. A further Phase 1 study exoASO-STAT6 (NCT05375604) aimed to assess the safety and preliminary antitumor activity of exosomes engineered to deliver an antisense oligonucleotide (ASO) targeting STAT6. While no data is available, the treatment aimed to reprogram macrophages from an immune-suppressive to an immune-stimulatory phenotype, potentially boosting the host antitumor response as observed in preclinical studies in hepatocellular and colorectal carcinoma [38]. Despite the exciting genesis of clinical trials and pre-clinical models there remains a paucity of data in relation to immune response to EVs in cancer. Understanding MSC-EV–immune interactions will be critical to successful translation.

This is the first study to assess the systemic administration of both human and murine MSC-EV on the host immune response in a model of breast cancer. Regardless of human or murine source no immune response was detected in any of the immune cell populations post EV administration. Although a small study, this is an important milestone in EV therapy and shows that MSC-EVs appear to retain the immune privilege of the MSCs. Human EVs will be employed in the clinical setting going forward and this data indicates that MSC-EVs may be tolerated well. Long-term studies with repeat dosing and increased number of EVs will be needed alongside investigation of use in combination with chemotherapy. After tumour resection there may be a rebounding of the immune system and EV administration post resection will be interesting to investigate [39]. There are many other aspects of EV biology that need to be further understood to support application in clinical oncology, including factors controlling migratory itinerary and recipient cell uptake, and the impact of genetic modification.

## 5. Conclusions

For translation to the clinical setting it is imperative that at an early stage in development of therapeutic EVs, researchers address and report the potential EV-host immune interactions. MSC-EVs offer immense potential in the therapeutic setting and may support development of “off the shelf” therapies that negate the need for cell recovery and expansion. In this study, EV administration resulted in no discernible detection by the host immune system, suggesting MSC-EVs retain the secretory cell properties. Further validation to show that MSC-EVs retain immune privilege in a long-term study and robust tracking of EV migratory itinerary and uptake is still required to confirm the homing properties. However, these exciting results reinforce the therapeutic potential of MSC-EVs.

## Acknowledgments

All flow cytometry experiments were performed in the University of Galway Flow Cytometry Core Facility, which is supported by funds from University of Galway, Science Foundation Ireland, the Irish Government ‘s Programme for Research in Third Level Institutions, Cycle 5, and the European Regional Development Fund. The authors wish to thank the Bio-Resources Unit technical, veterinary, and administrative staff in University of Galway for facilitating in vivo studies.

