## Supplementary Figures for "Investigation of immune response to Mesenchymal Stromal Cell-derived Extracellular Vesicles in the cancer setting"

### Supplementary Figure 1

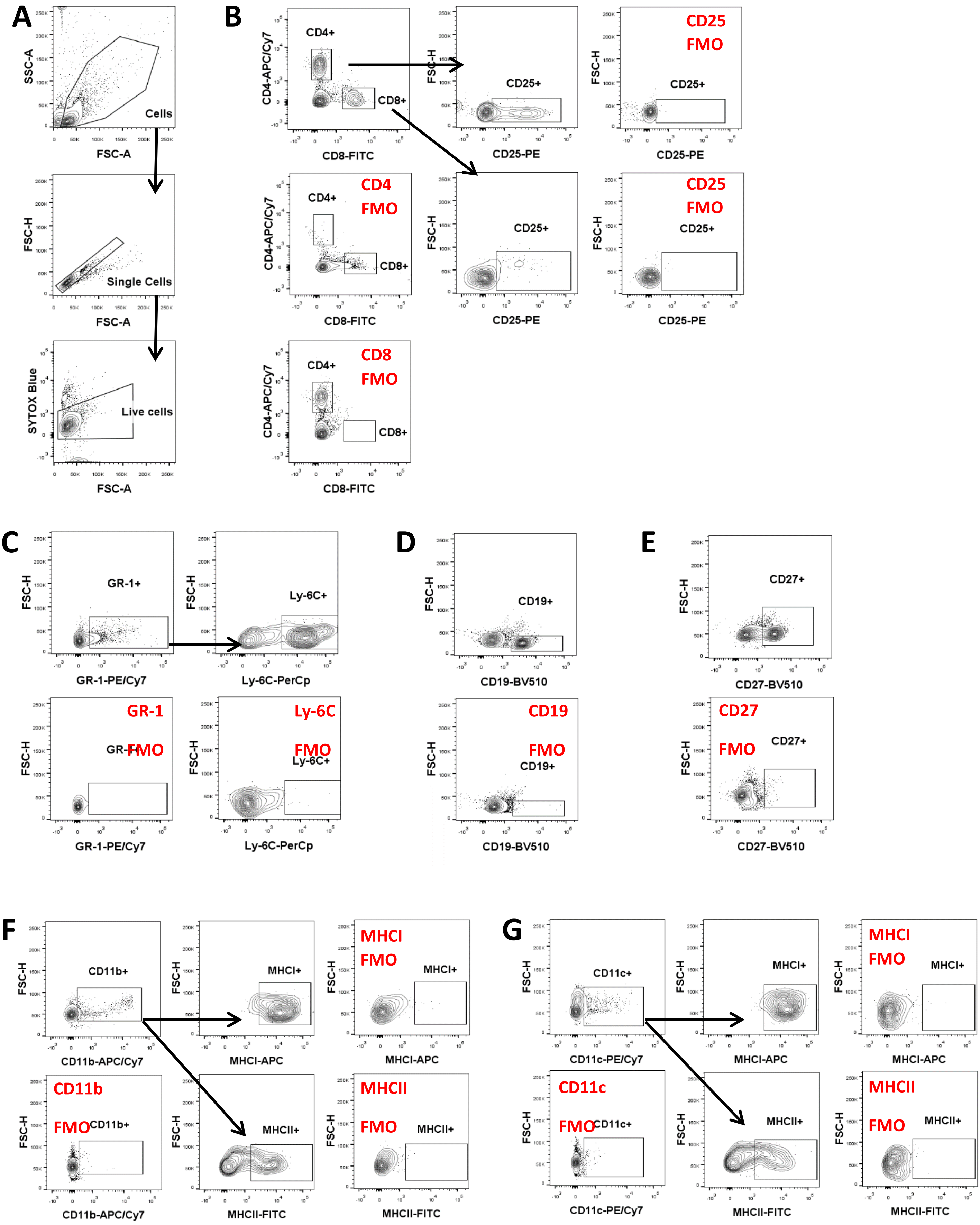

**Supplementary Figure 1**: Representative flow cytometry gating strategy used to select (A) cell population of interest → single cells → live cells, (B) activated T cells (CD4+CD25+ or CD8+CD25+), (C) neutrophils (GR-1+Ly-6C+), (D) B cells (CD19+), (E) NK cells (CD27+), (F) macrophages (CD11b+MHCI+MHCII+) or dendritic cells (CD11c+MHCI +MHCII+). The backbone gating shown in (A) was performed for all subsequent gating of cell populations shown in parts B-G. The gating shown here was performed on splenocytes and is representative of the gating used to select the same populations in lymph nodes and tumour-derived single cell suspensions.

### Supplementary Figure 2

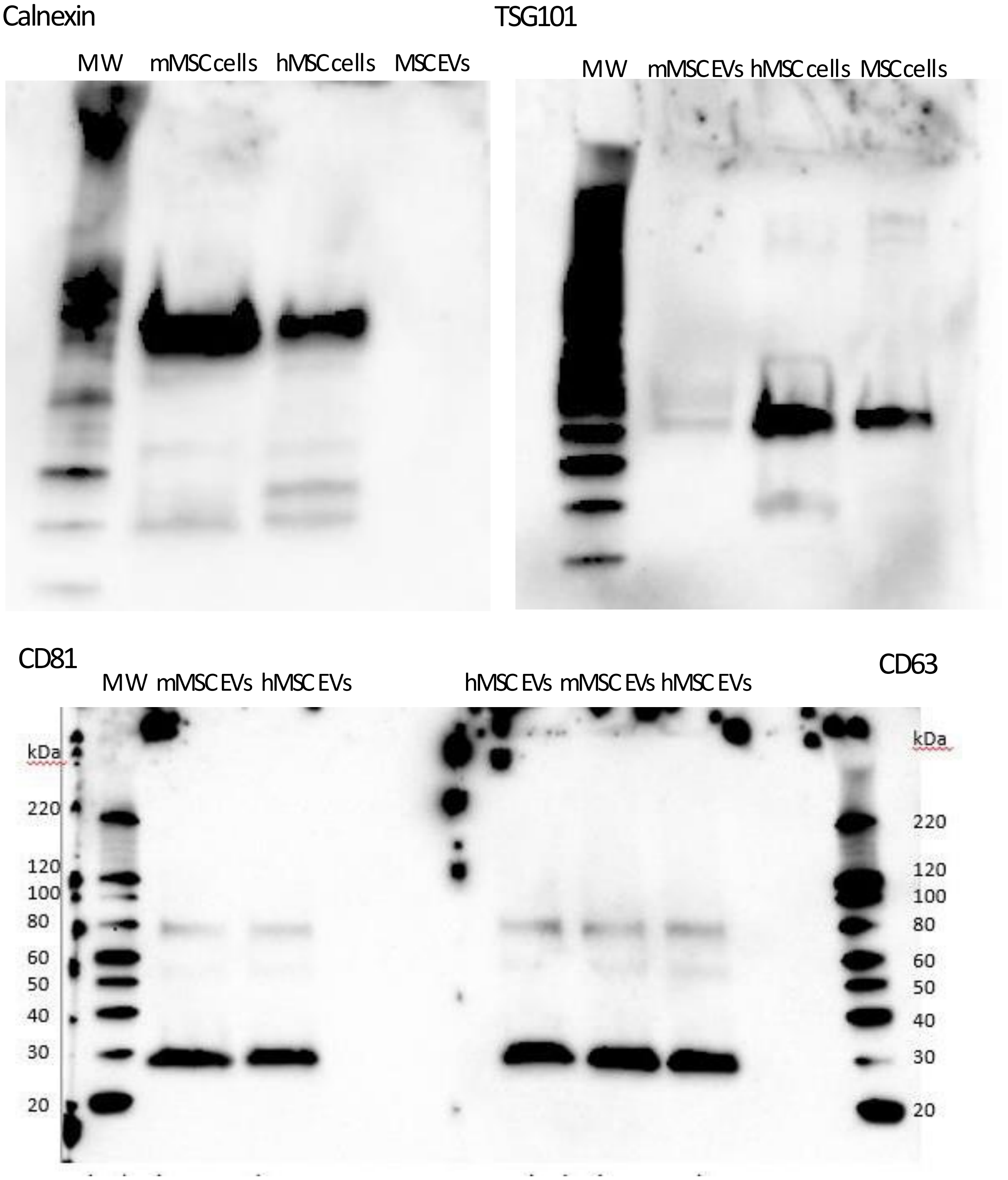


**Supplementary Figure 2:** Representative full face western blots showing the presence of CD81, CD63, TSG101 and the absence of Calnexin in MSC-EVs. MSC cell extracts are included as a positive control.
